# A microendovascular system can record precise neural signals from cortical and deep vessels with minimal invasiveness

**DOI:** 10.1101/2024.08.29.610418

**Authors:** Takamitsu Iwata, Hajime Nakamura, Takafumi Uemura, Teppei Araki, Takaki Matsumura, Takaaki Abe, Toshikazu Nezu, Masatoshi Takagaki, Tomohiko Ozaki, Shinpei Miura, Ryohei Fukuma, Sam E. John, David B. Grayden, Haruhiko Kishima, Tsuyoshi Sekitani, Takufumi Yanagisawa

## Abstract

Minimally invasive intravascular electroencephalography (ivEEG) signals are a promising tool for developing clinically feasible brain–computer interfaces (BCIs) that restore communication and motor functions in paralyzed patients. However, current ivEEG techniques can only record signals from the superior sagittal sinus (SSS), making it challenging to record motor responses related to hand and mouth movements from brain regions distant from the SSS, despite their critical role in BCIs. Here, using micro intravascular electrodes, ivEEGs recorded from the cortical or deep veins of eight pigs could measure cortical activities with greater signal power and better spatial resolution than those recording in the SSS, thus allowing mapping of the sensorimotor and visual functional areas. Additionally, electrical stimulation in the cortical vein between the micro intravascular electrodes induced muscle contractions contralateral to the stimulated area in five anesthetized pigs. These results demonstrate that ivEEG using micro intravascular electrodes is a promising tool for developing BCIs.

## Introduction

Less-invasive methods for recording intracranial electroencephalography (EEG) signals are essential for developing a clinically feasible brain–computer interface (BCI) that can restore communication and motor functions in patients with severe paralysis and monitor and consequently regulate various types of brain activity^1^. Previous studies have developed invasive BCIs that use various signal sources that achieve sufficiently high performance to support paralyzed patients, but less invasive methods are still needed^2^. In the early stages of invasive BCI development, devices that record multiunit activity (MUA) were important for achieving clinical feasibility and the ability to support precise motor reconstruction and reliable communication for patients^2^. The use of MUAs has made clinically feasible communication at high speeds with an error rate of less than 1% possible^3,4^. However, MUA-based devices have the problems of high invasiveness and low signal stability^5^. Therefore, approaches using intracranial EEG, such as electrocorticography (ECoG), have been investigated to achieve more stable and feasible BCIs^6^. In particular, the use of high-density microelectrocorticography (μECoG) along with neural decoding based on deep neural networks and natural language processing has improved the performance of communication BCIs^7,8^. One such speech BCI achieved a communication rate of 78 words per minute with comparable accuracy to that of MUAs^3,4,8^. In addition, a system using recordings with epidural electrodes enabled a patient with spinal cord injury to walk on his own legs by stimulating his spinal cord on the basis of the recorded signals^10^. Similarly, subdural and epidural electrodes implanted over the motor areas associated with the leg, hand and mouth have allowed some stable and feasible BCIs to be realized^11,12^. These advancements demonstrate that clinically feasible BCIs can be achieved with less invasive signals, such as those recorded by ECoG, through improvements in neural decoding.

Intravascular EEG (ivEEG) is a minimally invasive technique for recording intracranial EEG signals without craniotomy^13–15^. Previous studies have demonstrated that a stent attached with electrodes (called the “Stentrode™”, Synchron, USA) could be chronically implanted in the superior sagittal sinus (SSS) of sheep intravascularly to record intracranial EEG signals as well as subdural and epidural electrodes can^14–16^. In addition, the ivEEG was able to record somatosensory evoked potentials (SEPs) from the somatosensory cortex adjacent to the SSS^16,17^. It has also been demonstrated that an ivEEG recorded by Stentrode implanted in the SSS can be used to evaluate cortical activity related to foot movements stably in paralyzed patients and can be applied for BCI control of external devices^18,19^. However, ivEEG recordings using the Stentrode are currently limited to acquiring signals from the SSS, making it difficult to record motor cortical activity related to the hands and mouth, the main signal sources in current invasive BCIs^2–4,6–9^. To use ivEEG to record various cortical signals for use in the BCI, smaller intravascular electrodes (called micro intravascular electrodes) that can access the cortical veins and deep brain structures, such as the areas of the motor cortex related to the hand and mouth or visual cortex, are needed.

Furthermore, the electrodes in the SSS and cortex are separated by the dura matter and the subdural and subarachnoid spaces^20,21^. Shorter distances between recording electrodes and the cortex are required more precise and easily localized cortical signals^22^. Therefore, the ivEEG signals recorded from SSS may have a lower signal–to– noise ratio (SNR) than ECoG signals recorded by subdural electrodes on the cortical surface because of the distance from the cortex and the low-pass filtering due to the dura matter. The SNR of the ivEEG signals in SSS has been reported to be similar to that of the ECoG signals acquired with epidural electrodes^13, 16^. However, micro intravascular electrodes positioned in cortical veins are closer to the cortex because the vessel walls are thin, potentially allowing them to provide higher-quality cortical signals.

Here, we hypothesize that ivEEG recordings from electrodes in the cortical or deep veins can measure cortical activity with a higher SNR than those from electrodes in the SSS and identify areas related to motor and visual functions with higher spatial resolution. We used clinically available micro guide wires as electrodes and implanted them into the cortical vein and deep veins of both miniature and domestic pigs to compare the signal quality of ivEEG recorded in the SSS and to evaluate the efficacy of recording functional responses evoked by somatosensory and visual stimulation for mapping the functional areas of the sensorimotor and visual cortices.

## Results

### Endovascular approach to the intracranial veins of a miniature pig

We performed experiments on eight anesthetized pigs (five *Clawn* miniature pigs and three *Zen-noh Premium* pigs; Supplementary Table 1). For all pigs, a linear incision was made in the skin of the left groin area, and 5 Fr and 8 Fr Super arrow-flex sheaths (Teleflex Medical Japan, Tokyo, Japan) were placed in the left femoral artery and vein, respectively. From the 5 Fr Super arrow-flex sheath, a 4 Fr Cerulean (Medikit, Tokyo, JAPAN) was inserted into the internal carotid artery, and venography was performed (Fig. 1a, Supplementary Video 1). The diameter (mm) of each pig’s intracranial vessels is shown in Supplementary Table 1. From the 8 Fr Super arrow-flex sheath, an 8 Fr FUBUKI catheter (Asahi Intec, Aichi, JAPAN) was inserted into the cervical portion of the internal jugular vein. DeFrictor BULL (Medicos Hirata, Osaka, JAPAN) and Marathon (Medtronic, Minneapolis, MN, USA) microcatheters were then guided to the intracranial veins through the 8Fr FUBUKI catheter. An ASAHI CHIKAI 10 (Asahi Intec, Nagoya, Aichi, JAPAN) was placed in the transverse sinus (TS), SSS, cortical veins, or deep veins through the microcatheters (Fig. 1b, c; Supplementary Table 2). Notably, the tips of the guidewires for the SSS were slightly bent, with the curved part of the wire against the vessel walls.

**Figure 1.**
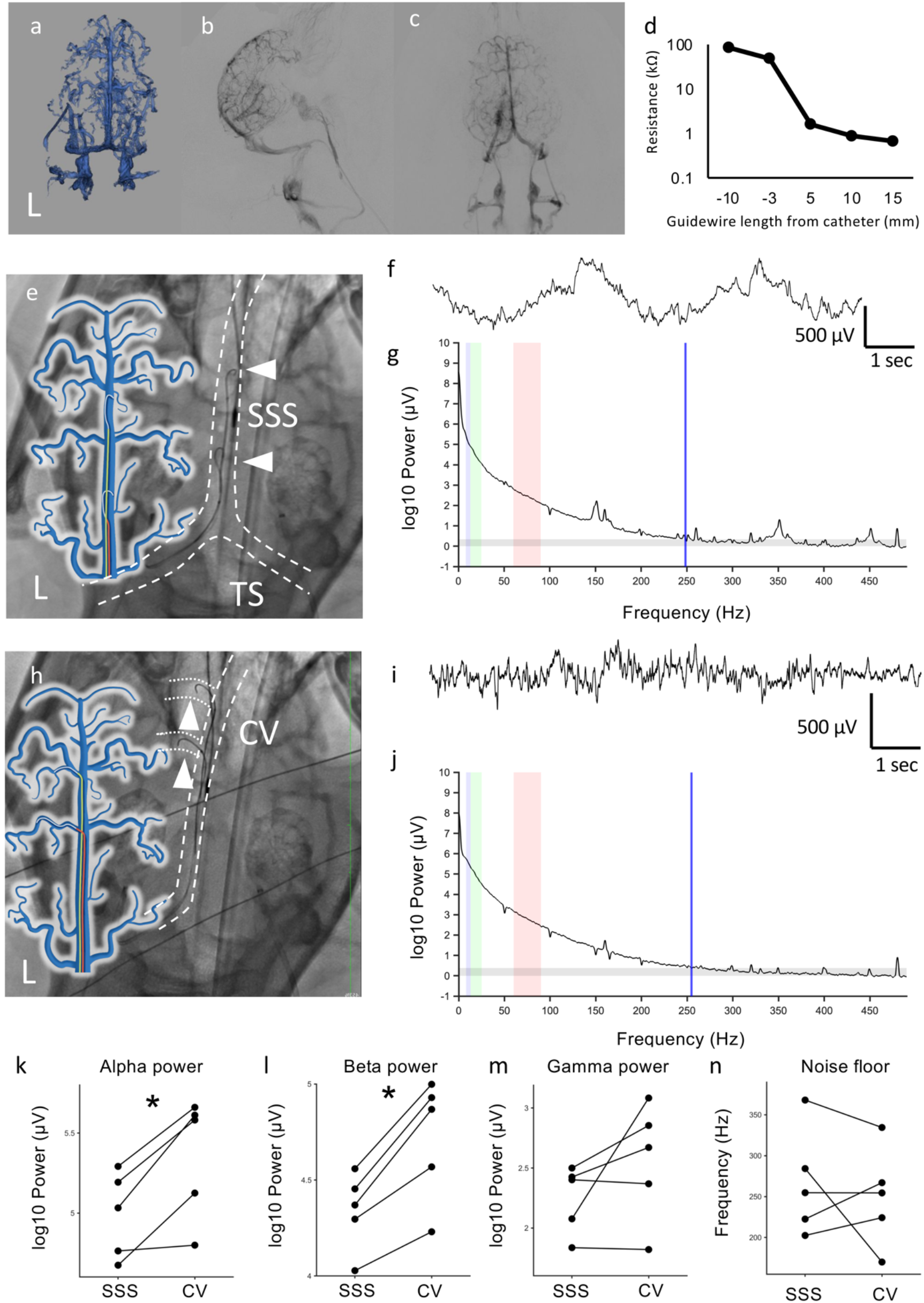
ivEEG measured with the micro-intravascular electrodes implanted in the SSS and cortical vein (a-c) Angiography of miniature Pig A; 3D reconstruction image (a) and subtraction image of the lateral view (b) and AP view (c) of the venous phase. (d) The impedance between two micro guide wires was assessed for various lengths of the guidewire beyond the catheter when the micro guidewires and catheter were inserted into Pig A. (e) Venography and illustration of the two guidewires placed in the SSS (white arrowheads) of Pig A. The dashed line shows the SSS and TS. (f) Representative waveforms of the CV-ivEEG. (g) Power spectrum density of the SSS-ivEEG. The blue, green and red shaded areas represent the alpha, beta and gamma power bands, respectively. The gray band indicates the noise level. The blue line indicates the noise floor. (h) Venography image and illustration of two guidewires placed in the left cortical veins (white arrowheads). The dashed line indicates the SSS and cortical veins. (i) Representative waveforms of the CV-ivEEG. (j) Power spectrum density of the CV-ivEEG. The blue, green and red areas represent the alpha, beta and gamma power bands, respectively. The gray band indicates the noise level. The blue line indicates the noise floor. (k-m) The PSDs of the SSS-ivEEG and CV-ivEEG data were compared for three frequency bands–alpha (k), beta (l) and gamma (m)–among five pigs. \**p*<0.05, paired t test, Bonferroni corrected. (n) Noise floors of SSS-ivEEG and CV-ivEEG for five pigs.

### Intravascular EEG of the SSS and CV

To record ivEEG signals, we removed the polymer coating on the tip and end of each micro guidewire (ASAHI CHIKAI 10 (Asahi Intec, Nagoya, Aichi, JAPAN)) and used the guidewires as micro intravascular electrodes. First, we assessed the impedance between the two guidewires in the SSS by changing the length of the guidewires extending beyond the insulated catheter (DeFrictor BULL (Medicos Hirata, Osaka, Japan)). When both guidewires were placed inside the catheter, the impedance was greater than 50 kΩ (Figure 1d); however, the impedance decreased to less than 1 kΩ when at least 5 mm of both guidewires was exposed beyond the tip of the catheter (Figure 1d). Similarly, in the saline solution, the impedance changed accordingly depending on the length of exposed guidewire from the catheter (Supplementary Table 3). Afterward, we recorded the intravascular EEG with the tip 5–10 mm from the catheter.

First, for Pig A, the ivEEG was measured between the micro intravascular electrodes in the SSS (Fig. 1e) and cortical vein (Fig. 1h) for 5 minutes without any stimulation. Stable ivEEG signals were recorded in the SSS (SSS-ivEEG, Fig. 1f) and cortical vein (CV-ivEEG, Fig. 1i) while the pig was anesthetized. The ivEEG recordings were mostly stable, but sometimes, the baseline exceeded the recording range. After these artifact segments were removed, the power spectrum densities (PSDs) of the ivEEGs were assessed (Fig. 1g, j; Methods). For both SSS-ivEEG and CV-ivEEG, the PSDs showed 1/f decay and exceeded the noise floor (based on the power in the 450--500 Hz range) up to the gamma range (SSS-ivEEG; 255 Hz, CV-ivEEG; 255 Hz). The PSDs of CV-ivEEG showed more signal power than those of SSS-ivEEG for Pig A (alpha: 52 dB (SSS-ivEEG), 56 dB (CV-ivEEG); beta: 46 dB (SSS-ivEEG), 50 dB (CV-ivEEG); and gamma: 25 dB (SSS-ivEEG), and 29 dB (CV-ivEEG)). Similarly, we recorded both CV-ivEEG and SSS-ivEEG for the other four pigs (Pigs B, C, D, and E). Among the five pigs, the PSDs showed significantly greater signal power for CV-ivEEG than for SSS-ivEEG for the alpha (8–13 Hz; *p*=0.0154, *t*(4)=-4.05, *n*=5, paired *t* test, Bonferroni corrected) and beta (13–25 Hz; *p*=0.0031, *t*(4)=-6.39, *n*=5) frequencies but not gamma frequencies(60–90 Hz; *p*= 0.1747, *t*(4)=-1.65, *n*=5) (Fig. 1k–m). Additionally, the noise floors of the SSS-ivEEG and CV-ivEEG recordings were not significantly different (mean: 266.4, 95% CI: 209.6–323.2 Hz for SSS-ivEEG; mean: 250.1, 95% CI: 197.3–302.8 Hz for CV-ivEEG; p>0.05; paired t test; n=5; Fig. 1n). Nonetheless, these results demonstrated that CV-ivEEG yielded larger signals for the alpha and beta ranges than did SSS-ivEEG.

### Somatosensory evoked potential (SEP)

To evaluate how the CV-ivEEG could identify localized functional activities, the somatosensory evoked potential (SEP) was assessed (Methods). First, we recorded SSS-ivEEG using two micro intravascular electrodes placed at the anterior and posterior regions of the SSS in Pig C. The tips of both micro intravascular electrodes in the SSS were placed against the left wall of the SSS to record the ivEEG of the left hemisphere. The SEPs of the right and left median nerves of the pigs’ forearms were subsequently compared (Fig. 2a). The N20 amplitude was not significantly different between the two stimulation sides (*p*>0.05, paired t test, Bonferroni corrected; Fig. 2b). Next, we recorded SEPs using CV-ivEEG from the micro intravascular electrodes placed in the left anterior cortical vein and the posterior cortical vein of Pig C (Fig. 2c). A clear N20 was observed in response to the stimulation of the right median nerve; additionally, the N20 amplitude was significantly greater than that for the stimulation of the left median nerve (*p*<0.05, paired t test, Bonferroni correction; Fig. 2d). Similarly, lateralized SEPs were observed for two other pigs (Pigs D and E) (Fig 2e-l). These results suggest that CV-ivEEG is more sensitive to the anatomical location of electrodes than SSS-ivEEG is and can be used for functional mapping of the somatosensory cortex.

**Figure 2.**
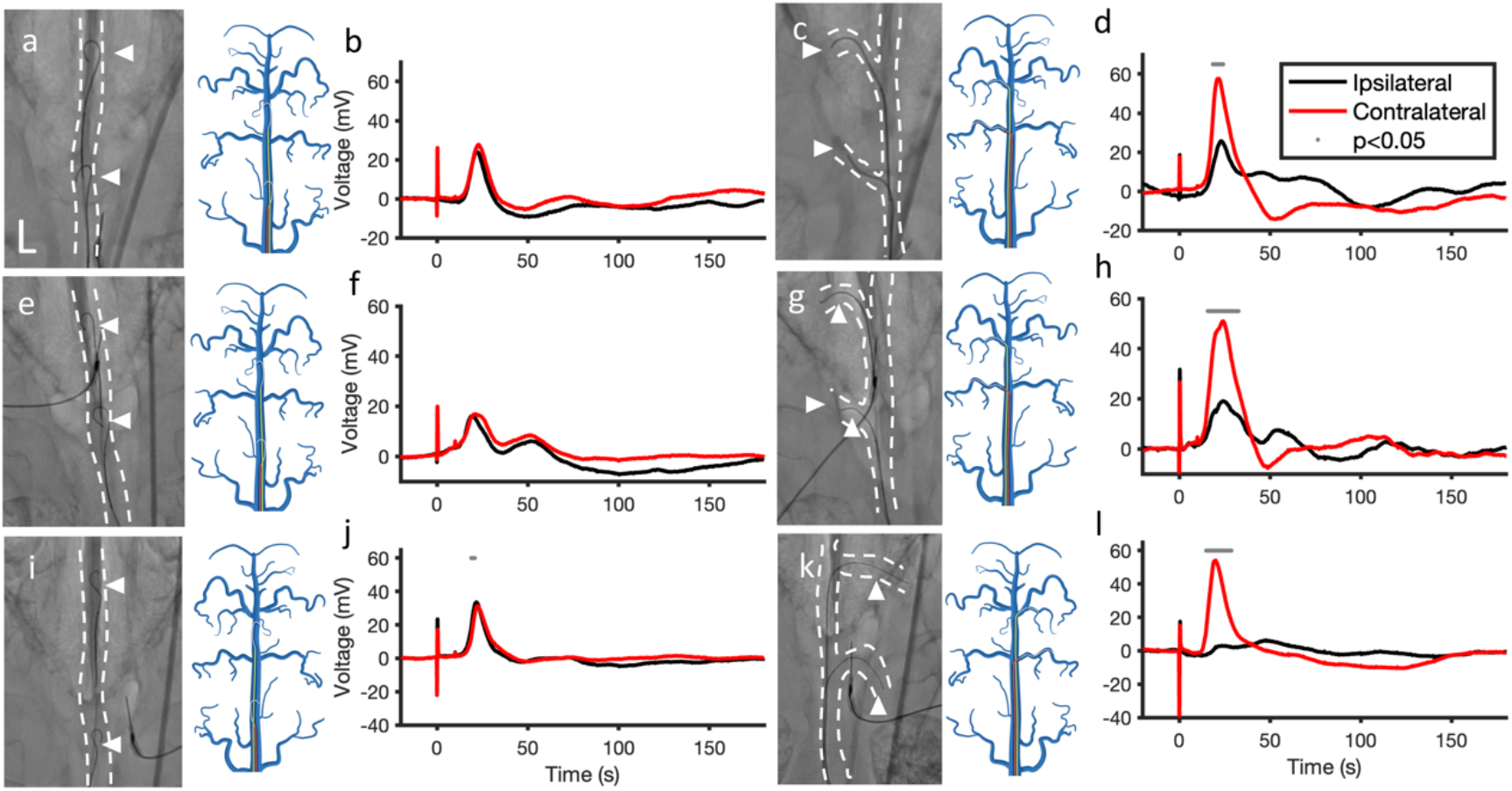
(a) Venography and illustration of SSS-ivEEG in Pig C. The electrodes were attached to the right wall of the SSS. (b) SEP recorded by SSS-ivEEG in Pig C. The red and black lines correspond to the responses to stimulation of the contralateral (left) and ipsilateral (right) forearms of the pig. (c) Venography image and illustration of CV-ivEEG in Pig C. The electrodes were placed in the left cortical veins. (d) SEP recorded by CV-ivEEG in Pig C. The red and black lines correspond to the response stimulated to the contralateral (right) and ipsilateral (left) forearms of the pig. The gray dots indicate the times at which the amplitudes of contralateral and ipsilateral stimulation were significantly different (*p*<0.05, paired t test, Bonferroni corrected). (e) Venography and illustration of SSS-ivEEG of Pig D. (f) SEP of SSS-ivEEG in Pig D. (g) Venography and illustration of CV-ivEEG of Pig D. (f) SEP of CV-ivEEG in Pig D. (i) Venography and illustration of SSS-ivEEG of Pig E. (f) SEP of SSS-ivEEG in Pig E. (g) Venography and illustration of CV-ivEEG of Pig E. (f) SEP of CV-ivEEG in Pig E.

To compare the CV-ivEEG data with the ECoG data, we implanted epidural and subdural electrodes into two pigs (Pigs C and D) to collect CV-ivEEG data. For Pig C, we implanted epidural electrodes on the left frontal lobe close to the CV electrodes (Fig. 3a). Conducting simultaneous CV-ivEEG and epidural ECoG recordings without stimulation revealed that the CV-ivEEG had a PSD similar to that of the epidural ECoG (Fig. 3b). In addition, recording simultaneous with somatosensory stimulation revealed that the laterality of the SEP amplitude was greater in the CV-ivEEG recordings than in the epidural ECoG recordings (Fig. 3c, d). Similarly, we implanted subdural electrodes close to the CV electrodes in Pig D (Fig. 3e). The PSD of recordings during the resting state demonstrated that gamma power was significantly greater in the ECoG recordings than in the CV-ivEEG recordings (Fig. 3f). On the other hand, the laterality of SEPs from CV-ivEEG and ECoG recordings was similar (Fig. 3g, h), although the amplitude of CV-ivEEG SEPs was greater than ECoG SEPs. These results demonstrate that CV-ivEEG records cortical responses with similar signal power to those of the ECoG and sometimes assesses functional localization better than ECoG recordings do.

**Figure 3.**
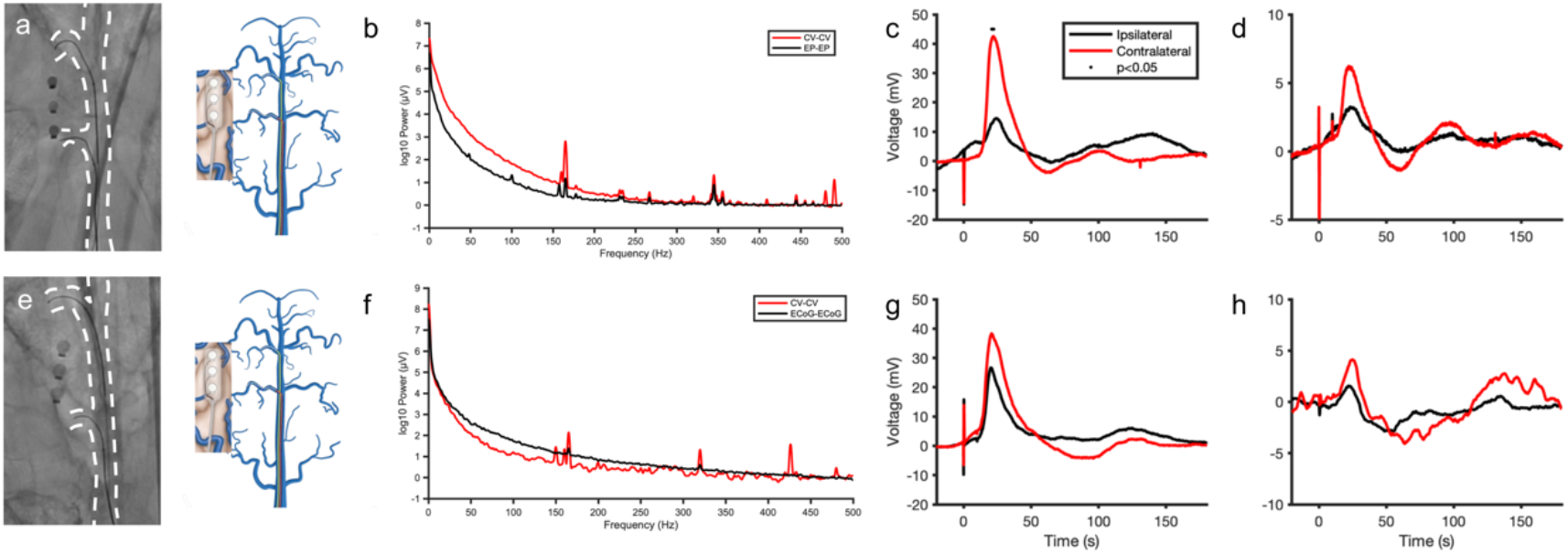
Simultaneous recording of CV-ivEEG and ECoG (a) X-ray image and illustration of implanted electrodes of Pig C. (b) PSDs of CV-ivEEG and epidural ECoG during the resting state. (c, d) SEPs resulting from stimulation of the forearms contralateral and ipsilateral to the implanted electrodes in the (c) CV and (d) epidural spaces. The black dots indicate time points when the amplitudes of contralateral and ipsilateral stimulation were significantly different (*p*<0.05, paired t test, Bonferroni corrected). (e) X-ray image and illustration of implanted electrodes of Pig D. (f) PSDs of CV-ivEEG and subdural ECoG during the resting state. (g, h) SEPs resulting from stimulation of the forearms contralateral and ipsilateral to the implanted electrodes in the (g) CV and (h) subdural space.

To further assess functional localization based on the phase reversal of SEPs, we implanted three guidewires in the SSS and cortical veins of two pigs (Pigs F and G) (Fig. 4a, c). The N20 of the SEP was compared between two adjacent guidewires. The phase reversal of the N20 was confirmed for both pigs (Fig. 4b, d). These results indicate that ivEEG modalities, including CV-ivEEG, can map the sensorimotor cortex by measuring the phase reversal of SEPs.

**Figure 4.**
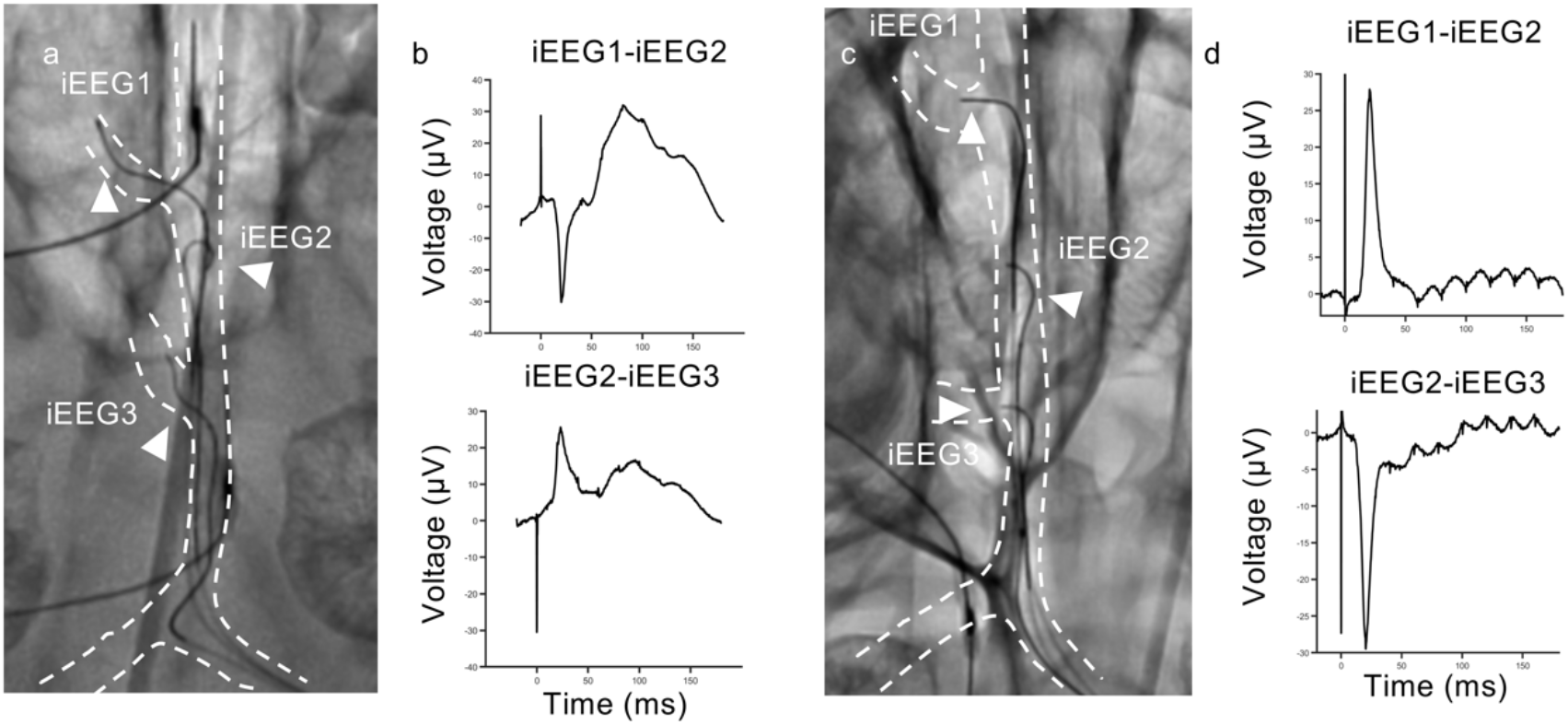
Phase reversal observed by ivEEGs (a) X-ray of the micro- and intravascular electrodes of pig G (b) SEPs from stimulation of the forearms contralateral (red) and ipsilateral (black) to the implanted electrodes (ivEEG1-ivEEG2 and ivEEG2-ivEEG3). (c) X-ray of the micro- and intravascular electrodes of pig F. (b) SEPs resulting from stimulation of the forearms contralateral to the implanted electrodes (ivEEG1-ivEEG2 and ivEEG2-ivEEG3).

### Motor evoked potentials (MEP) measured through the cortical intravascular system

For five pigs (Pigs A, B, C, F, and H), electrical stimulation was applied through the micro intravascular electrode to identify the motor cortical responses. The muscle responses were assessed by measuring motor-evoked potentials with electrodes inserted into the muscles of the pigs’ lips and shoulders and confirmed by observation of muscle contraction. For Pigs A and B, reproducible localized muscle responses to electrical stimulation with electrodes implanted in the SSS were not observed. For Pig A, electrical stimulation with a biphasic pulse of 50 Hz between the electrodes at the left anterior CV and SSS induced muscle contraction of the right shoulder (Fig. 5a). On the other hand, the same stimulation at the left anterior CV and left posterior CV induced muscle contraction of the right upper lip (Fig. 5b). In addition, the short electrical stimulation of 5 biphasic pulses of 500 Hz in the same electrode pairs induced a clear motor-evoked response recorded by the electrodes at the right upper lip (Fig. 5c, Supplementary Video 2). Similarly, among the other four pigs, cortical stimulation of the micro intravascular electrode induced muscle twitches on the contralateral face and shoulder (Fig. 5d-h). Notably, in Pig H, stimulation of the electrodes at the frontal pole induced no muscle contraction, although stimulation of the middle part of the brain induced a contralateral muscle contraction. These findings suggest that CV electrodes can identify the motor cortex by assessing localized MEP responses.

**Figure 5.**
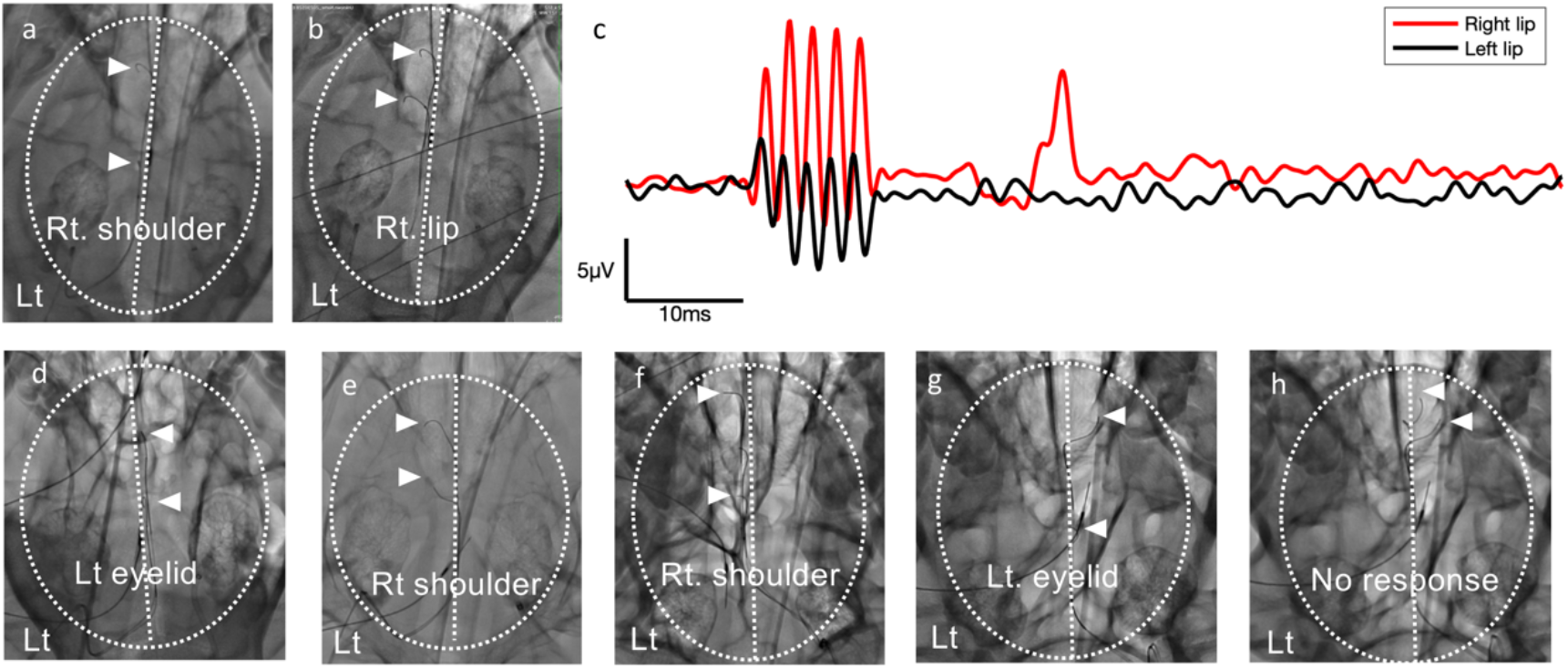
Cortical stimulation to induce motor responses (a) X-ray image of Pig A showing the location of the electrodes (white arrowheads). The white dotted line shows the skull and center line of the pig’s head. Stimulation at 50 Hz on the electrodes induced muscle contractions on the right shoulder of the pig. (b) Electrode location of Pig A. Electrical stimulation at 500 Hz induced muscle contraction of the right lip. (c) Electromyogram recorded from the right (red) and left (black) sides of the lip of the pig. (d) Electrical stimulation at 50 Hz (white arrowheads) in Pig B induced muscle contractions of the left side of the face. (e) Electrical stimulation at 50 Hz in Pig C induced movements of the right shoulder. (f) Stimulation at 50 Hz on Pig F induced right shoulder movements. (g) Stimulation at 50 Hz in Pig H induced movements of the left eye lid. (h) Stimulation at 50 Hz induced no muscle contraction in Pig H.

### Visual evoked potentials measured through deep veins

Visual evoked potentials (VEPs) were recorded via ivEEG from the deep vein. ivEEG signals were recorded between two guidewires placed in the internal cerebral vein (ICV) and TS (Fig. 6a). In addition, ivEEG was recorded between the micro-intravascular electrode at TS and the scalp electrode attached at the back of the pig’s head (Oz). The following light stimulation was applied to both eyes: intensity 2000 lux, frequency 3 Hz and duration 20 msec. Then, the VEP was measured. The results revealed a clear peak potential of approximately 40 msec in both cases, but the amplitude of the intravascular EEG was greater with the ICV-TS than with the Oz-TS (Fig. 6b and 6c).

**Figure 6.**
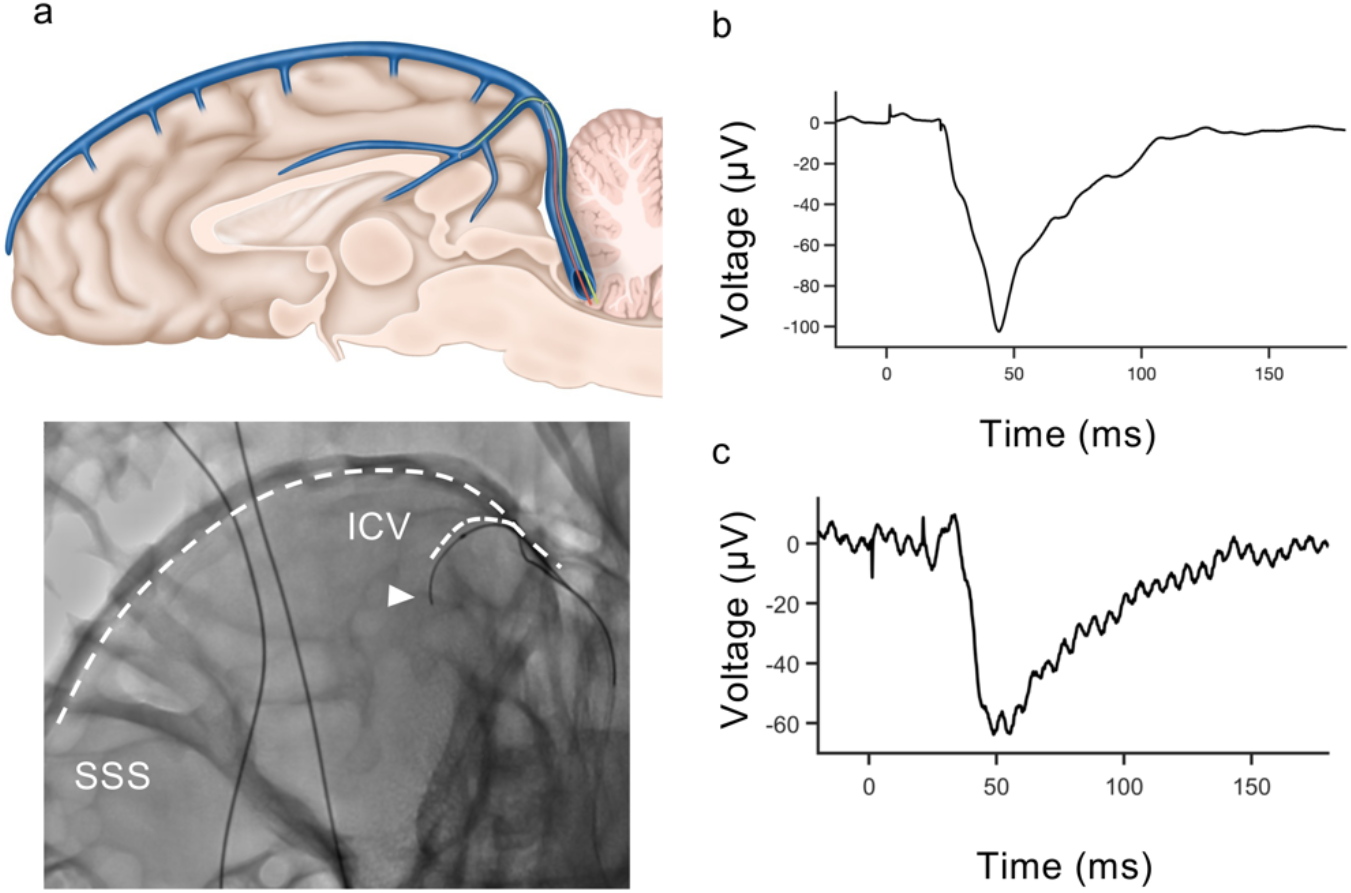
Visual evoked potentials recorded by ivEEG in the ICV (a) Illustration (upper panel) and X-ray image (lower panel) of the implanted electrodes in Pig F. (b, c) VEP recorded by micro intravascular electrodes between the (b) ICV and TS and between the (c) TS and Oz.

### Safety assessment of intravascular cortical electrodes

We placed micro guidewires (CHIKAI10) safely into the cortical and deep veins of eight pigs. During the insertion of the micro guidewires, we spent 5–6 hours recording ivEEG data while frequently repositioning the electrodes. However, we observed no obvious extravasation or any other finding that might suggest vascular injury. In addition, some pigs with micro guidewires implanted in the CV and SSS were craniotomized, but we did not observe any obvious vascular injuries, such as subarachnoid hemorrhage, indicating that micro guidewires can be safely used to record ivEEG data from the cortical and deep veins. In general, the vessel diameters of miniature pigs are small: 1.9 (0.5) mm for the SSS, 0.5 (0.29) mm for the CV and 1.4 (0.2) mm for the internal jugular vein (Supplementary Table 2). Notably, the size of the SSS in miniature pigs corresponds to the size of the cortical vein in humans^23^. These results in miniature pigs suggest that the microendovascular system can be used to safely access the cortical vein in the human brain and measure intravascular EEGs.

## Discussion

This study demonstrated that ivEEG recorded by micro guidewires (micro-intravascular electrode) in the cortical vein and deep vein can measure functional cortical responses with amplitudes that are comparable and sometimes larger than those of ECoG and that various cortical regions can be accessed for functional mapping by this less invasive tool. Compared with those of SSS-ivEEG, the CV-ivEEG PSDs demonstrated greater signal power in the alpha and beta ranges. Compared with those measured with SSS-ivEEG, the SEPs measured by the CV-ivEEG demonstrated greater contrast to the stimulated side than those measured with SSS-ivEEG, suggesting that CV-ivEEG can detect spatially localized responses. Using micro-intravascular electrodes, both the MEP and the phase reversal of the N20 could be identified, facilitating mapping of the motor cortex. In addition, VEPs could be measured from the ICV or SSS via micro intravascular electrodes, demonstrating high accessibility to various brain regions that are difficult to record with subdural electrodes. These results demonstrated that ivEEG can yield high SNRs and area-specific recordings of intracranial EEGs with minimal invasiveness by using a electrode to access the cortical and deep veins.

Our results suggested that CV-ivEEG has a greater SNR than SSS-ivEEG both in terms of the PSD and SEPs. In addition, the SNR of CV-ivEEG was comparable to that of epidural and subdural ECoG. These results are consistent with previous studies showing that SSS-ivEEG has a similar SNR to epidural ECoG and a smaller SNR than subdural ECoG^16^. The SNR of EEG is affected by both the distance of the electrode from the cortex and the intervening tissue; in the SSS, the dura mater and cerebral fluid act as low-pass filters, attenuating high-frequency band activity. Because the superficial cerebral vein has thin vascular walls and is close to the cortex, CV-ivEEG provides a higher SNR signal than SSS-ivEEG does. Therefore, implanting electrodes in the CV seems to provide a higher SNR EEG compared to EEG measured in the SSS.

Using a micro guidewire to access cortical and deep vessels, cortical region-specific signals can be obtained. For intravascular EEG in the SSS, it was also possible to measure hemispheric-specific SEPs depending on whether the electrode was placed on the left or right wall. However, obtaining localized muscle responses with electrical stimulation with electrodes implanted in the SSS was difficult. Nonetheless, electrical stimulation from wires in the CVs was shown to allow cortical area-specific motor cortical stimulation. In particular, the mouth region of pigs could be selectively stimulated, suggesting that the CV can be used to measure the motor cortex activity of the mouth, which is important for speech BCIs^7–9^. Notably, a clear correspondence between pigs and humans is difficult to establish because of differences in the functional arrangement in the sensorimotor cortex^24,25^. VEPs were also measured in the deep blood vessels. Visual responses can also be used for communication BCIs^26^. It has been shown that function-specific activity can be obtained with high accuracy by measuring ivEEG with a micro guidewire. Furthermore, this finding suggests the signals can be acquired from deep structures such as the brainstem and basal ganglia as well as previously reported methods allowing superficial regions to be recorded via the SSS and TS.

This study has several limitations. Since this study was conducted as an acute-phase experiment, we were unable to assess the long-term stability and safety of CV-ivEEG. However, no major adverse events occurred during the experiments. Additionally, in this study, CHIKAI10 was used for CV-ivEEG measurements; however, CHIKAI10 was not originally intended for this application. While the resistance values of the electrodes were sufficient for the EEG recordings, future work should focus on the development of dedicated electrodes that impose minimal strain on cortical veins while achieving high measurement precision.

## Conclusion

In this study, micro-intravascular electrodes were implanted in cortical and deep veins, demonstrating that micro-intravascular EEG has a high SNR comparable to that of ECoG and a higher spatial resolution for mapping sensorimotor and visual cortices that of SSS-ivEEG.

## Methods

### Study Design and Approval

All surgical and experimental procedures were performed at the Fukushima Medical Device Development and Support Center. The research protocol was approved by the Institutional Review Board (approval no. 04-073-000) and included an evaluation of catheter approaches to the brain surface and deep veins in pigs, an evaluation of micro guidewire-based EEG measurements from within the vessels, and a comparison of the acquired EEG signals with those acquired by ECoG. EEG quality assessment was performed by measuring SEPs, MEPs, and VEPs.

### Animal and experimental information

The smallest number of animals required to demonstrate feasibility was used in accordance with the policy of the institutional animal ethics committee.

Eight pigs (16.5 months old, 43.1 kg) were subjected to experimental procedures. (Supplementary Table 1)

The procedure was performed under general anesthesia, which was induced with isoflurane. Subsequently, general anesthesia was maintained with continuous infusion of propofol [2 mg/kg/h] and dexmedetomidine [0.5 mg/kg/h]). Additional doses of propofol or dexmedetomidine were administered as needed when body movements occurred.

The intracranial vascular anatomy of pigs is similar to that of humans, with the cortical vein flowing into the SSS and merging with the straight sinus from the inferior sagittal sinus (ISS). However, it differs from humans in that the transverse sinus drains into the vertebral venous plexus and internal jugular venous system via the sigmoid sinus.

### micro-intravascular electrode placement

Cerebral angiography was performed at the Fukushima Medical Device Development and Support Center. Groin puncture was performed with a cut-down procedure under ultrasound evaluation. 5 Fr and 8 Fr Super arrow-flex sheaths (Teleflex Medical Japan, Tokyo, Japan) were inserted into the femoral artery and vein, respectively. Heparin was administered at a dose of 100 international units (IU)/kg via an intravenous catheter placed in the ear vein, and an activated coagulation time longer than 250 seconds was maintained during the procedure. Digital subtraction angiography (DSA) was performed with an INFX-8000C (Canon Medical Systems, Tochigi, Japan); specifically, iopamidol was injected 3 ml at 0.5 ml/sec as contrast medium via a 4 Fr Cerulean catheter placed in the internal carotid artery. Additionally, 3-D rotational angiography/venography was performed to evaluate the precise vascular architecture, with 5 ml at 0.5 ml/sec injection of contrast medium.

To access cortical veins from the femoral vein, an 8 Fr FUBUKI catheter (Asahi Intec, Aichi, JAPAN) was introduced into the internal carotid vein via 8 Fr Super arrow-flex sheaths by using coaxial catheter and a 0.035 inch guidewire. The microcatheters were maneuvered into the intracranial veins (sigmoid sinus, transverse sinus, superior sagittal sinus, internal cerebral vein, and cortical veins) through the jugular bulb by using 0.010-inch guidewire. In this procedure, roadmap navigation under the ideal projection was needed to clearly identify the venous route.

After the microcatheter reached the target vessels, a 0.010-inch guidewire was pulled back, and then the ivEEG electrodes were positioned at the target vessels using the microcatheters. In the venous sinus, the 5 mm tip of the wire was bent into a J shape to make contact with the vessel wall. However, the diameter of the cortical veins was approximately 1 mm; therefore, high-definition angiographic equipment and delicate catheter guidance techniques were essential, and the tip was not bent in the cortical veins.

### EEG measurement

Scalp and intracranial EEGs were measured using a Power Lab 4/26 with an FE232 Dual Bio AMP (ADInstruments, Dunedin, New Zealand). The recordings were sampled at 10000 Hz. When intracranial EEG was measured, an alligator clip was attached to the tip of the CHIKAI 10 and connected to the EEG system. The end-to-end resistance of CHIKAI 10 was sufficiently low, approximately 40 Ω. EEG recordings were obtained without applying a low-pass filter, but a high-pass filter with a cutoff frequency of 0.1 Hz was used. To remove the influence of body movement due to breathing, the wires of the recording electrodes were fixed to the bed or the floor. The ground electrode was placed on the skin of the neck or shoulder.

The Fz electrode was inserted into the nasal root, and the Cz electrode was inserted into the midline of the cranium; placement was confirmed in the frontal fluoroscopy image. The impedance measurements were made at a frequency of 100 Hz via an LCR meter IM3590 (HIOKI E.E. Co., Nagano, JAPAN) or LCR-916 (GW Instek, TAIWAN).

When the impedance was measured, the distance the guidewire protruded from the catheter tip was visually assessed on the radiographic fluoroscopic image.

### Power spectral density

The recorded signals were segmented into 2-s epochs, and epochs with signals exceeding the recording limits were excluded from further analysis. To remove power line noise, a 50 Hz notch filter and 0.5-500 Hz bandpass filter were applied. The noise of each electrode was estimated using the median spectral power between 455 Hz and 495 Hz of each recording electrode. This frequency band was chosen because it is the highest frequency band located below the Nyquist frequency and does not include any harmonics of the 50-Hz line noise. The noise floor was defined as the frequency at which the power first dropped below the upper limit of noise (mean + 1 SD). Since ECoG power spectra are not normally distributed and show considerable variability within subjects, the mean power spectrum was normalized to the mean noise power and log-transformed.

### SEP

SEPs were measured in a cohort of five pigs using microivEEG. SEPs were recorded under general anesthesia. A modular stimulator NS-101 (UNIQUE MEDICAL, Tokyo, Japan) was used to generate a stimulation current. Needle electrodes were inserted into the median nerve of the bilateral forelegs for bipolar electrical stimulation^27,28^. Stimulations with an amplitude of 25–30 mA, a pulse width of 0.2 s, and a stimulus frequency of 4.3 Hz were applied and repeated 200 times. The stimulation amplitude was changed for each experimental condition. Neural signals were bandpass filtered with a 0.1 Hz high-pass filter. Averaged SEP waveforms were normalized by the mean of the baseline signals (from -20 ms to 0 ms).

### MEP

A alligator clip was attached to the tip of the CHIKAI 10, and a modular stimulator NS-101 (UNIQUE MEDICAL, Tokyo, Japan) was used for stimulation. Stimulation was conducted at a frequency of 50 Hz, with a 20 mA current, a pulse width of 0.5 milliseconds and a duration of 3 seconds. Muscle contraction was visually detected before stimulation at 500 Hz (20 mA, pulse width of 1 millisecond, biphasic, in 5 trains)^27^. In this case, the MEP was measured by inserting a needle electrode into the shoulder and upper lip muscles using a Power Lab 4/26 system with a FE232 Dual Bio AMP (ADInstruments, Dunedin, New Zealand).

### VEP

Stimulations were conducted using an LED light stimulation device (LFS-101 III Unique Medical, Tokyo, Japan) in a lighting room. LED pads were applied over the eyelids of both closed eyes. The stimulation conditions were 20,000 lux, 20 msec stimulation duration, 3 Hz stimulation frequency, and 200 repetitions.^29^

### ECoG recording

The electrocorticography electrodes consisted of three pole electrodes mounted on a silicon sheet measuring 5 mm by 15 mm (Unique Medical, Tokyo, Japan). Each electrode had 3 mm diameter contacts and electrodes were placed 5 mm apart. The pigs were placed in a prone position for the placement of the electrocorticography electrodes. The appropriate position for craniotomy was determined via X-ray fluoroscopy. Then, a skin incision was made, part of the skull was removed using a drill, and the dura mater was exposed. A linear incision was carefully made in the dura mater, allowing for the placement of an electrocorticography electrode directly on the surface of the brain.

## Acknowledgements

This research was supported by the Japan Science and Technology Agency (JST) Moonshot R&D (JPMJMS2012).

## Author contributions

Conceptualization: T.Y., T.S., H.N.; methodology: H.N., T.S., T.U., T.A., R.F. and T.Y.; investigation, data curation: T.I. H.N., T.U., T.A., T.M., T.Ab., T.N., M.T., T.O., S.M., R.F., T.S., T.Y.; formal analysis, and software: T.I., T.Y., R.F.; funding acquisition: T.Y., H.N., T.S.; writing–original draft: I.T., H.N., T.U., T.Y.; writing–review and editing: all authors; supervision: T.Y., T.S.

## Additional information

Supplementary videos are available for this paper.

