## Supplementary information for "A microendovascular system can record precise neural signals from cortical and deep vessels with minimal invasiveness"

**This PDF file includes the following:**

Supplementary Table. 1 to 3

**Other Supplementary Materials for this manuscript include the following:**

Supplementary Movies 1 and 2

| Subject ID | Age (month) | Breed | Body weight (kg) | Position | Sheath placement |
|---|---|---|---|---|---|
| A | 28 | *Clawn miniature pig* | 47.6 | Supine | R. FA/R. FV |
| B | 22 | *Clawn miniature pig* | 41.8 | Prone | L. FA/L. FV |
| C | 3 | *Zen-noh Premium pig* | 38.8 | Prone | L. FA/L. FV |
| D | 3 | *Zen-noh Premium pig* | 44.4 | Prone | L. FA/L. FV |
| E | 3 | *Zen-noh Premium pig* | 45.0 | Prone | L. FA/L. FV |
| F | 23 | *Clawn miniature pig* | 43.6 | Prone | L. FA/L. FV |
| G | 25 | *Clawn miniature pig* | 41.2 | Prone | L. FA/L. FV |
| H | 25 | *Clawn miniature pig* | 42.6 | Prone | L. FA/L. FV |

**Supplementary Table 1**

Information on the pigs.

R. FA; right femoral artery, R. FV; right femoral vein, L. FA; left femoral artery, L. FV; left femoral vein.

| | IJV | | SS | | TS | | SSS | | | Cortical vein | | | | | |
|---|---|---|---|---|---|---|---|---|---|---|---|---|---|---|---|
| | Lt | Rt | Lt | Rt | Lt | Rt | Prox | Mid | Dist | Lt.Ant | Rt.Ant | Lt.Mid | Rt.Mid | Lt.Post | Rt.Post |
| A | 1.1 | NaN | 4.3 | 2.9 | 2.4 | 2.1 | 2.2 | 1.7 | 1.5 | 1.2 | 0.7 | 0.8 | 1..2 | 1 | NaN |
| B | NaN | NaN | 4.4 | 3.6 | 2.4 | 1.8 | 3 | 1.7 | 2 | 0.6 | 1.1 | 1.1 | 1.2 | 0.7 | 0.8 |
| C | 1.4 | 1.5 | 2.9 | 3.2 | 2.4 | 2.8 | 3.2 | 2 | 1.5 | 0.7 | 0.9 | 0.6 | 0.9 | 1.1 | 0.5 |
| D | 1.6 | 1.5 | 2.7 | 5 | 3.2 | 2.6 | 2.3 | 1.6 | 1.6 | 1.1 | 0.6 | 0.8 | 0.8 | 1.5 | NaN |
| E | 1.3 | NaN | 3.6 | 3.3 | 1.6 | 2 | 1.4 | 1.5 | 1.7 | 1.6 | 0.7 | 0.8 | 0.6 | 1 | NaN |
| F | NaN | 1.3 | 2.5 | 4 | 1.3 | 3.2 | 3 | 2.6 | 1.4 | 1.2 | 0.7 | 1.2 | 1 | 0.6 | NaN |
| G | 1.2 | 1.4 | 2.8 | 3.1 | 1.7 | 2 | 1.6 | 1.5 | 1.5 | 1.3 | 0.9 | 0.6 | 0.7 | NaN | 1.1 |
| H | NaN | 1.5 | 2.5 | 3.6 | 2.7 | 2.8 | 2.1 | 1.6 | 1.4 | 1 | 1.4 | 1 | 1.5 | 1.5 | 0.7 |
| Mean (SD) | 1.4 (0.2) | | 3.4 (0.7) | | 2.3 (0.6) | | 1.9 (0.5) | | | 0.95 (0.29) | | | | | |

**Supplementary Table 2**

The diameter (mm) of each pig's intracranial vessels.

First, the cortex was divided into three segments: anterior, middle, and posterior. The diameter of the cortical vein with the largest diameter in each segment was measured. The diameter of the superior sagittal sinus (SSS) was measured at three specific points in the frontal view: just after the confluence of the sinuses (proximal), just distal to the middle cortical vein (middle), and just distal to the anterior cortical vein (distal). The diameter of the transverse sinus (TS) was measured at the midpoint between the confluence of the sinuses and the transverse-sigmoid junction. Additionally, the diameter of the straight sinus (SS) was measured at the point where the shape was linear, with the smallest variation in diameter.

IJV: internal jugular vein, TS: transverse sinus, StS: straight sinus, ICV: internal cerebral vein, SSS: superior sagittal sinus, Lt: left, Rt: right, Ant: anterior, Mid: middle, Post: posterior.

| | Distance from catheter tip [mm] | +20 | +15 | +10 | +5 | 0 | -3 | -5 | -10 |
|---|---|---|---|---|---|---|---|---|---|
| 10 Hz | Z (kΩ) | 0.43 | 0.55 | 0.77 | 1.46 | 14.37 | 22.69 | 37.23 | 68.27 |
| | θ(°) | -44.51 | -44.62 | -44.89 | -44.97 | -7.06 | -2.44 | -2.09 | -1.99 |
| 100 Hz | Z (kΩ) | 0.16 | 0.19 | 0.26 | 0.47 | 12.32 | 21.20 | 35.39 | 65.40 |
| | θ(°) | -32.25 | -33.75 | -36.96 | -41.78 | -4.15 | -1.82 | -1.47 | -1.93 |

**Supplementary Table 3**
Distance of the CHIKAI 10 Exposure from the Defirector nanobull Tip and the corresponding impedance in a saline solution.
